# Metabolic silencing via methionine-based amino acid restriction in multiple myeloma cell lines reveals a potential new strategy for cancer therapy

**DOI:** 10.1101/2025.08.17.670755

**Authors:** Andrea Frabschka, Maximilian F. Völter, Hannah Jöhren, Werner Schmitz, Charlotte Wiegand, Mohamed El-Mesery, Corinna Koderer, Lena Wiesner, Alexander Christian Kübler, Axel Seher

**Affiliations:** Department of Oral and Maxillofacial Plastic Surgery, University Hospital Wuerzburg, Wuerzburg, Germany; Department of Biochemistry and Molecular Biology, Biocentre, Wuerzburg, Germany; Department of Biochemistry, Faculty of Pharmacy, Mansoura University, Mansoura, Egypt

## Abstract

Nutrient restriction, from caloric to amino acid to protein restriction, promotes cell switching to low-energy metabolism (LEM), which is characterized primarily by inhibited cell proliferation. In the long term, nutrient restriction has enormous impacts on life span and age-associated diseases such as type 2 diabetes and cancer. Owing to their antiproliferative effects, nutrient restriction approaches with a focus on amino acids are receiving increasing attention for use in tumour therapy. However, to date, the effects of amino acid restriction (AAR) have rarely been reported in patients with multiple myeloma (MM). In this work, we analysed AAR implemented via methionine restriction (MetR) in the MPC11 murine cell line and two human cell lines, KMS12-BM and L363. MetR exerted profound antiproliferative effects without causing cell death. In addition, methionine dependency was demonstrated in the cell lines investigated via homocysteine compensation. Additionally, mass spectrometry analyses of MPC11 cells revealed metabolic reprogramming, indicating that MetR led to a dramatic decrease in the (aerobic) glycolysis rate and in the abundances of energy equivalents and nucleotide synthesis metabolites. Taken together, the results of this study suggest that MetR is an important therapeutic option for MM.

## Introduction

Multiple myeloma (MM) is a B-cell lymphoma characterized by monoclonal proliferation of plasma cells in the bone marrow, leading to a large increase in secreted immunoglobulins in the form of Para- or Bence Jones proteins and ultimately to plasmacytosis. People with MM typically show increased susceptibility to infections and organ damage, which may involve massive destruction of bone structures (osteolysis) [1–3]. The incidence of MM is approximately four to six new cases/100,000 people per year. This rate corresponds to 10% of all haematological cancers or 1% of all cancers [4]. Many drugs are available to treat MM, in addition to alkylators and corticosteroids. Thalidomide, lenalidomide, and pomalidomide are immunomodulatory agents. Bortezomib, carfilzomib, and ixazomib are proteasome inhibitors. Elotuzumab and daratumumab are monoclonal antibodies that target the surface antigens CD319 (SLAMF7) and CD38, respectively. Finally, panobinostat is a deacetylase inhibitor [5]. In recent years, chimeric antigen receptor (CAR)-T cells have been increasingly used as a therapeutic option [6]. The survival of patients with MM has been prolonged significantly in the past 20 years [7]. However, the 5-year survival rate of 55.6% is still too low [8], even though the median overall survival in younger patients (aged <50 years) was >10 years by 2014 [9]. An expansion of the possible therapeutic options is clearly desirable.

The most pronounced feature of neoplastic cells is their unlimited proliferation [10]. Two factors are needed for cell division: energy and mass. While the energy level of a cell is reflected by ATP and NAD(P)H equivalents, amino acids constitute the majority of the cellular mass [11].

For this reason, the components necessary for cell proliferation are measured at the molecular level through a sophisticated strategy: ATP is measured by AMP kinase [12], NADH is measured by sirtuins [13], and selected amino acids (e.g., leucine, arginine, glutamine (Gln), serine, and methionine) are measured on the basis of different protein complexes [14].

The amino acid methionine is measured indirectly on the basis of the SAMTOR complex, which is formed by the intermediate S-adenosylmethionine (SAM) [15]. Many of the measured signals converge at a central complex, mechanistic target of rapamycin (mTOR), that is evaluated to determine whether a cell is proliferating or not proliferating [16]. When the energy and mass levels are sufficient, proliferation can be initiated. However, a lack of even one component inhibits proliferation, and autophagy is usually induced. In this state, which we call low-energy metabolism (LEM), the cell is not in a state of starvation; therefore, cell viability is not compromised. In contrast, the cell both reorganizes its own genome through the action of sirtuins and recycles valuable metabolites through induced autophagy [17,18].

LEM can be induced by various forms of nutrient restriction. Caloric restriction (CR), amino acid restriction (AAR) and protein restriction (PR) are the most common forms. All of these forms of nutrient restriction induce LEM. When applied in the long term, these different restriction regimens lead to profound effects. The complete life span is extended, and age-associated diseases, such as cardiovascular diseases, type 2 diabetes and cancer, are prevented. Equally remarkable, these effects can be achieved across organismal kingdoms, since a large portion of the effect is based on mechanisms conserved throughout evolution [19–21].

Within the past decade, the benefit of nutrient restriction in cancer therapy has become increasingly clear. One reason for the effectiveness of nutrient restriction is the ability to induce them medicinally through caloric restriction mimetics (CRMs). Well-known examples of CRMs are aspirin and metformin, which are currently being used successfully in cancer therapy and clinical trials [22–24].

In this work, we analysed, for the first time, the possible use of AAR as a therapy for MM. We implemented an AAR strategy targeting the amino acid methionine, which is particularly suitable for this treatment because of its key role in metabolism. Methionine restriction (MetR) is one of the oldest, most common, and most successfully used forms of AAR. The effectiveness of MetR has been proven through numerous studies, e.g., research in the field of cancer [25–27]. The central position of methionine in both metabolism and protein synthesis predestined this amino acid for restriction-based therapy [26,28]. In addition, many tumour cells are methionine dependent, losing the ability to regenerate methionine from homocysteine (Hcy) [26,29], as proven in 1979 with a cell line of acute lymphoblastic leukaemia (CCRF-HSB-2 cell line), among other cell lines [30]. This dependence on methionine is exacerbated in cancer cells, which show an increased need for methionine, as indicated by neoplastic cells “stealing” methionine from the environment or depriving surrounding tissues and cells of this resource by increasing their expression of the membrane methionine transporters SLC7A5 and SLC43A2, among other tactics [31,32].

The strong inhibitory potential of MetR was demonstrated in this work via proliferation assays in the murine cell line MPC11 and the human cell lines KMS12-BM and L363. All three cell lines were found to be methionine dependent, as Hcy-mediated compensation was impossible.

In our laboratory, the MPC11 murine cell line served as a model system for MetR. For proof of principle, liquid chromatography/mass spectrometry (LC–MS) was performed to generate a metabolic profile (metabolic fingerprint) of MetR every 24 h over a five-day period. More than 170 metabolites derived from amino acids through glycolysis and the tricarboxylic acid (TCA) and urea cycles and from purines, pyrimidines, etc., were examined to demonstrate the effects of MetR at the metabolic level. This work fundamentally demonstrates the potential of AAR as a new strategy for MM treatment.

## Materials and methods

### Cell culture

The MPC11 murine MM cell line was purchased from the American Type Culture Collection (ATCC), and the KMS12-BM and L363 human MM cell lines were purchased from the Leibniz Institute, DSMZ-German Collection of Microorganisms and Cell Cultures GmbH (Braunschweig, Germany). The MPC11 cells were cultured in DMEM, and the KMS12-BM and L363 cells were cultured in RPMI 1640 medium (both from Gibco, Life Technologies; Darmstadt, Germany) supplemented with 10% foetal calf serum (FCS; Sigma–Aldrich, Darmstadt, Germany) and 1% penicillin/streptomycin (P/S; 100 U/mL penicillin and 100 µg/mL streptomycin; Thermo Fisher Scientific, Darmstadt, Germany) at 37 °C in a humidified atmosphere containing 5% CO_2_. The L929 murine fibroblast line was purchased from the Leibniz Institute, DSMZ German Collection of Microorganisms and Cell Cultures GmbH (Braunschweig, Germany). These cells were cultured in RPMI 1640 medium (Gibco, Life Technologies; Darmstadt, Germany) supplemented with 10% FCS (Sigma–Aldrich, Darmstadt, Germany) or 1% P/S (100 U/mL penicillin and 100 μg/mL streptomycin (Thermo Fisher Scientific, Darmstadt, Germany)) at 37 °C in a humidified atmosphere containing 5% CO_2_.

For the AAR experiments, the following media, all lacking the amino acid methionine, were used. For the controls (complete medium), the amino acid methionine (Sigma–Aldrich, Darmstadt, Germany) was added at the indicated concentrations. To prepare the Met(-) medium, no amino acids were added. The DMEM (Gibco, Life Technologies; Darmstadt, Germany) contained 30 mg/L L-methionine. The RPMI 1640 medium (Gibco, Life Technologies; Darmstadt, Germany) contained 15 mg/L L-methionine. For the compensation experiments, in every case, 1.5 µM vitamin B12 and 800 µM D-/L-Hcy, 800 µM D-/L-Hse or 800 µM SAM were added.

### ImageXpress pico automated cell imaging system - digital microscopy (pico assay)

The cells were seeded at 10,000 cells in 100 µL of culture medium per well in a 96-well plate and incubated in complete or methionine-free medium in triplicate. The incubation time is stated in the corresponding figure legend. For staining, 10 µL of Hoechst staining solution (1:200 dilution in Hoechst 33342 medium (10 mg/mL in H_2_O); Thermo Fisher, Darmstadt, Germany) was added to each well, and the samples were analysed after 20–30 min of incubation. The wells were analysed with an ImageXpress Pico automated cell imaging system (Molecular Devices, San Jose, CA, USA) via automated digital microscopy. The cells were analysed with transmitted light and in the DAPI channel at 4× magnification. The complete area of every well was screened. The focus and exposure time were set via an automated setup and controlled by analysing 3–4 test wells. Finally, every result was confirmed visually, and 95% or more of the cells in the population were counted and analysed.

### Analysis of the cell progression rate via the pico assay

The cells were seeded at 10,000 cells in 100 µL of culture medium per well in a 96-well plate. After 0, 6, 24, 30, 48, 54, 72, 78 and 96 h, cell numbers were quantified, with six values for every time point as described in the Pico Assay Section (2.2). The growth of a cell population can be described with the following formula:

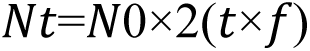

(Nt = cell number at time t; N0 = cell number at time 0; t = time in days (d); f = cell division frequency (1/d)).

To determine f, the formula is rearranged as follows:

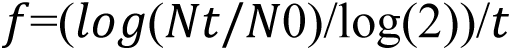

To obtain an overview, the measured values were first plotted as a simple diagram. From this, it was possible to determine at what point the growth entered the plateau phase. From these values, the individual f values were calculated for the intermediate periods (e.g., Δ24/30, Δ30/48). The total value f was then calculated as the mean of the Δf values.

### Live/dead cell viability assay

In a 96-well plate, 100 µl of a cell suspension with 100,000 cells/ml was seeded in complete or Met(-)-containing medium. As a death control, one well from each experiment was stimulated with 1 µM staurosporine (Seleckchem, Planegg, Germany). Measurements were performed after 24 h, 48 h and 72 h with an EarlyTox Live/Dead Assay Kit (Molecular Devices). A staining solution of 1 ml of DMEM, 5 µl of 10 mg/ml Hoechst 33342, 15 µl of 4 mM calcein AM and 30 µl of 2 mM ethidium homodimer-III (EthD-III) was prepared, and 10 µl of this staining solution was added to each well for measurement. After 30 min of incubation, measurements were performed with the “Cell Scoring: 3 Channels” program of the ImageXpress Pico Automated Cell Imaging System (Molecular Devices). To determine the absolute cell number, measurements were performed in transmitted light and in the DAPI channel via Hoechst 33342, as well as via the FITC (CAM) and Texas Red (TRITC)/(EthD-III) channels over the entire area of the respective well at 4× magnification. The exposure time and focus were determined by the automated settings. For better visualization of the cells, the measurement was subsequently repeated with a selected area of the well (1.39 mm × 1.39 mm) at 10× magnification, from which the images shown in this publication were selected. For each experimental condition, two wells were evaluated. The experiment was performed two times.

### MPC11 cell experiments for LC‒MS

A total of 2×10^6^ MPC11 cells were seeded in 25 mL of medium in 15 cm Petri dishes, and every value was measured on the basis of a 30 mg/L L-methionine control and 0.5 mg/L L-methionine for Met(-) and incubated for 24 h, 48 h, 72 h, 96 h and 120 h. The cells were removed from the Petri dishes with a cell scraper. The pellets were produced via centrifugation at 1500xg for 5 min. The cells were resuspended in the appropriate medium, and the cell number was determined in every sample at four different times to obtain an accurate number of cells. A total of 1×10^6^ cells were transferred to a 1.5-mL Eppendorf tube and centrifuged at 18000xg for 5 min. The supernatant was removed, and until LC–MS analysis, the pellets were stored at −20 °C. The experiment was performed three times.

### LC–MS

After the addition of 0.5 mL of MeOH/CH_3_CN/H_2_O (50/30/20, v/v/v) containing 10 µM lamivudine, the cell pellets were homogenized via ultrasonication (10*1 s at 250 W output energy). The external standard lamivudine was not used for absolute metabolite quantification but was used as a quality control to compensate for inevitable technical issues. For quality control and determination of the corresponding retention times, most of the annotated metabolites (which are commercially available) were run as mixtures of pure compounds under identical experimental conditions. General procedure: The resulting suspension was centrifuged (20 kRCF for 2 min in an Eppendorf centrifuge 5424), and the supernatant was loaded onto a C18-SPE column that was activated with 0.5 mL of CH_3_CN and equilibrated with 0.5 mL of MeOH/CH_3_CN/H_2_O (50/30/20, v/v/v). The solid-phase eluate (SPE) was evaporated under vacuum. The resulting pellet was dissolved in 50 µL of cell extract in 5 mM NH_4_OAc in CH_3_CN/(25%/75%, v/v).

Mobile phase A consisted of 5 mM NH_4_OAc in CH_3_CN/H_2_O (5/95, v/v), and mobile phase B consisted of 5 mM NH_4_OAc in CH_3_CN/H_2_O (95/5, v/v).

After the sample was loaded onto a ZIC-HILIC column (at 30 °C), the following LC gradient programme was applied: 100% solvent B for 2 min, a linear decrease to 40% solvent B within 16 min, maintenance at 40% solvent B for 9 min, and an increase to 100% solvent B within 1 min. The column was maintained at 100% solvent B for 5 min for column equilibration before each injection. The flow rate was maintained at 200 μL/min. The eluent was directed to the electrospray ionization (ESI) source of a Q Exactive (QE)-mass spectrometer from 1.85 min to 20.0 min after sample injection.

The MS parameters were as follows: a full MS scan was obtained in alternating positive- and-negative mode; the scan range was 69–1000 m/z; the resolution was 70000; the AGC target was 3E6; the maximum injection time was 200 ms; the sheath gas volume was 30; the auxiliary gas volume was 10; the sweep gas volume was 3; the spray voltage was 3.6 kV (in positive mode) or 2.5 kV (in negative mode); the capillary temperature was 320 °C; the S-lens RF level was 55.0; and the auxiliary gas heater temperature was 120 °C. Annotation and data evaluation: Peaks corresponding to the calculated monoisotopic masses (MIM +/- H+ ± 2 mMU) were integrated via TraceFinder software (Thermo Scientific, Bremen, Germany). Materials: Ultrapure water was obtained with a Millipore water purification system (Milli-Q Merck Millipore, Darmstadt, Germany). The high-performance LC (HPLC_–MS solvents, NH4OAc for use in LC–MS and lamivudine were purchased from Merck (Darmstadt, Germany). The RP18-SPE columns consisted of 50 mg of Strata C18-E (55 µm) in 1-mL tubes (Phenomenex, Aschaffenburg, Germany). A Branson Sonifier Ultrasonics 250 instrument equipped with a 13-mm sonotrode (Thermo Scientific, Bremen, Germany) was used.

A Thermo Scientific Dionex UltiMate 3000 ultra-HPLC (UHPLC) system linked to a QE-mass spectrometer equipped with a heated ESI (HESI) probe (Thermo Scientific, Bremen, Germany) was used. The samples were analysed with a high-resolution mass spectrometer, allowing the generation of extracted ion chromatogram (XIC) data that were then analysed by applying a very narrow m/z margin (+/- 3 mMU). A Javelin particle filter with an internal diameter (ID) of 2.1 mm (Thermo Scientific, Bremen, Germany) was used. The UHPLC precolumn was a SeQuant ZIC-HILIC column (5-μm particle size, 20 × 2 mm; Merck, Darmstadt, Germany). The UHPLC column used was a SeQuant ZIC-HILIC column (3.5-μm particle size, 100 × 2.1 mm; Merck, Darmstadt, Germany).

LC–MS analyses were carried out in three independent experiments at 24 h, 48 h, 72 h, 96 h and 120 h, with each value obtained from triplicate measurements. Metabolites were quantified in cell pellets under methionine-supplemented and methionine-reduced conditions (12 samples per time point in total). The resulting peak areas were normalized against that of lamivudine, which was the external standard. Hence, the mean value and standard deviation were calculated for each triplicate. For better comparisons, the values were converted to percentages. For the cell pellets, the first measured control value (24 h) in each test series within an experiment was defined as 100%. From these values, the average mean values from the two experiments were summarized and are presented in the individual tables. To create a better overview, the results were rounded to natural numbers and are shown in a heatmap. The corresponding colour ranges are indicated individually under each table. The raw data and results for the profiles are presented as Excel files in Supporting Files S3.

### Statistical analysis

Data collection and plotting were performed with Excel (Microsoft, Redmond, WA, USA) and GraphPad Prism (version 6.04; GraphPad Software, San Diego, CA, USA) software. Statistical analyses were performed with GraphPad. Comparisons between different groups were performed via one-way analysis of variance (ANOVA) followed by the Tukey–Kramer multiple comparison test.

## Results

Since MetR is essentially characterized as having a strong antiproliferative effect, cell proliferation rates were initially determined from f-values, which indicate how often a cell doubles in a period of 24 h. A value of 1 means that the cell doubles once per day, and a value of 0.5 means that the cell takes 48 h to double. Since many dynamic and limiting factors influence cell culture, we calculated a total of three different f values [33]. f-total indicates the division rate over the period examined until cell division enters the saturation phase. f-start indicates the division rate at the beginning of the experiment within the first 6 h, and f-max is the highest f-value and thus the maximum rate that was achieved within the measurement period. The results are shown in Fig 1. In contrast to adherent cells, MM cells in suspension require a certain phase of acclimatization during which the f-start values are correspondingly low. In the exponential growth phase, the values for all the cell lines are greater than one, with the murine cell line MPC11 having the highest division rate, with a value of 1.18. Owing to the slow start phase, the values of the three cell lines examined are well below 1: 0.78 for the murine cell line (MPC11) and 0.67 (L363) and 0.64 (KMS12-BM) for the two human cell lines.

**Fig 1.**
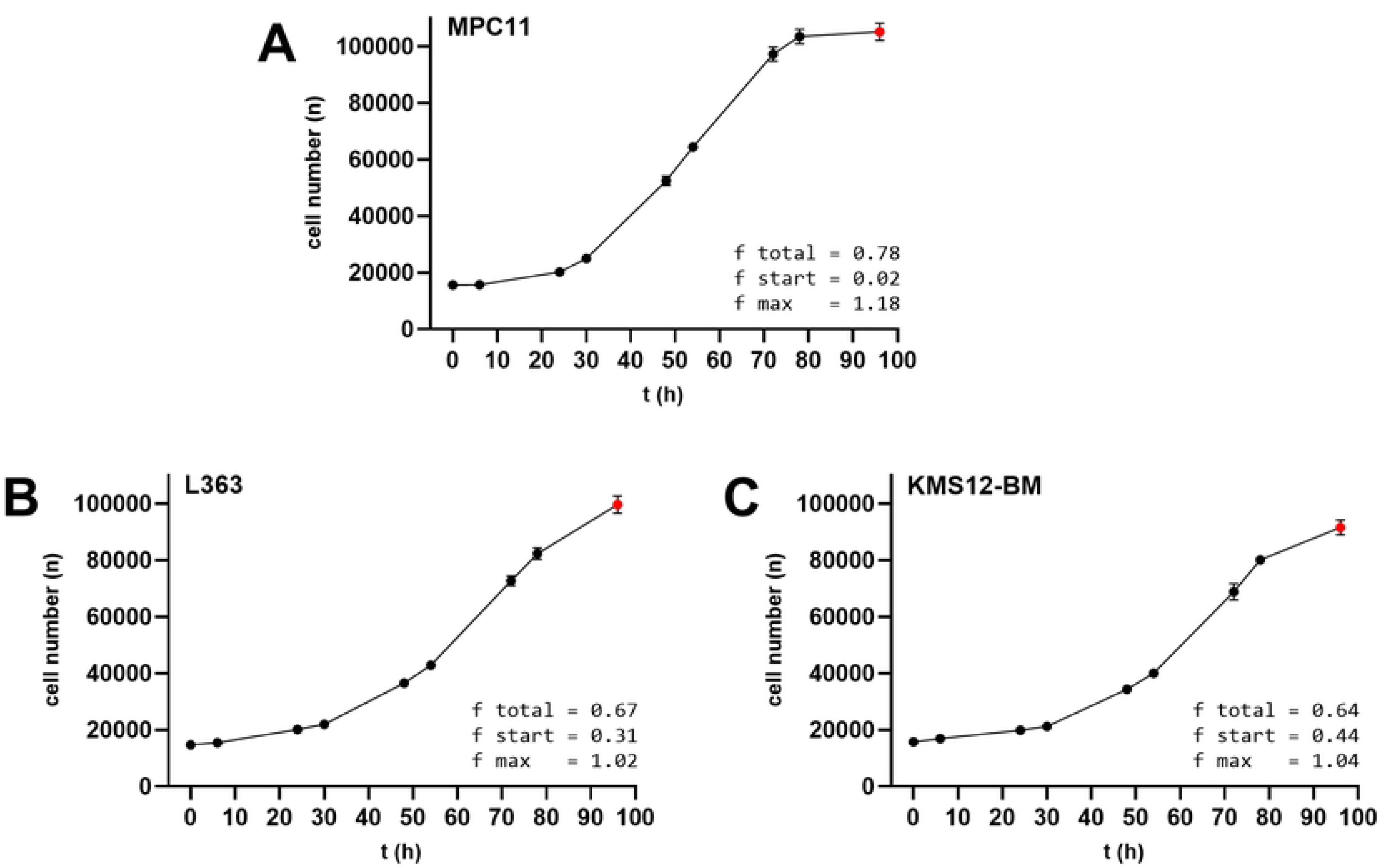
Analysis of the Cell Proliferation Rate. The cells were seeded at 10,000 cells/well in a 96-well plate. After 0, 6, 24, 30, 48, 54, 72, 78 and 96 h the cell numbers were measured with the ImageXpress Pico Automated Cell Imaging System as described in the Materials and Methods. The summarized results (n = 3) are shown. The values used for the analysis are marked in black, and the excluded measured values are marked in red.

### Methionine-based AAR exerts a strong antiproliferative effect on MM cells

In the next set of experiments, the potential of MetR to inhibit MM cell line proliferation was analysed. For this purpose, digital microscopy was performed because it enables automated counting of cells after the cell nucleus is stained. The advantage of this method is that the absolute number of cells at each well and point in time can be determined. Therefore, after the initial number of cells at the 0-h time point is determined, the efficiency of the inhibitory action can be accurately assessed, and the number of dead cells can be discerned. Nutrient restriction led to very efficient inhibition of MPC11 cell proliferation, which was highly significant after 48 h (Fig 2A). In general, the number of cells never reached a value greater than the initial number. MetR also inhibited proliferation very efficiently in the two human cell lines, L363 and KMS12-BM (Fig 2C and E). After 48 h, a significant difference in proliferative behaviour was already evident. In contrast to MPC11, the values under MetR were slightly increased compared with the initial value (t0).

**Fig 2.**
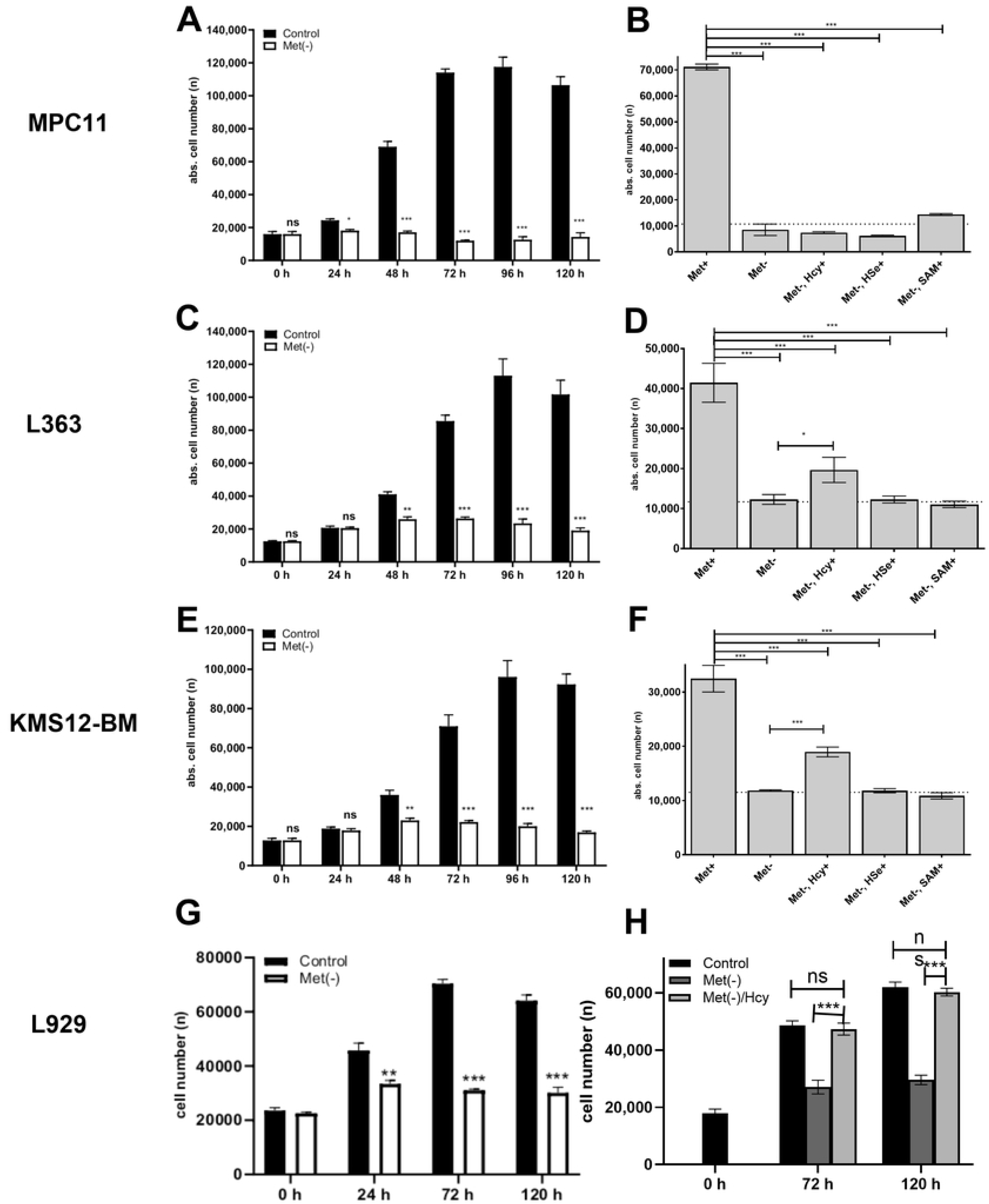
Analysis of Methionine Restriction (MetR) and Homocysteine (Hcy) Competition on Multiple Myeloma (MM) Cell Proliferation. (A, C, E) A total of 10,000 cells were seeded in each well and stimulated in triplicate with Met or without methionine (Met-). The proliferation of the cells was analysed through ImageXpress digital microscopy at the indicated time points as described in the Materials and Methods. The summarized results (n=3) are shown. (B, D, F) For Hcy-based competition with MetR, a total of 10,000 cells/well were seeded in 96-well plates and stimulated in triplicate. For the compensation experiments, 800 μM D-/L-Hcy D-/L-homoserine (Hse) or S-adenosylmethionine (SAM) was added to the Met(-)-medium. The cells were analysed via ImageXpress digital microscopy after 72 h, as described in the Materials and Methods. Comparisons between different groups were performed via one-way analysis of variance (ANOVA) followed by the Tukey–Kramer multiple comparison test. (ns, nonsignificant; ** p < 0.01 and *** p < 0.001). For better comparison, the previously published results for the adherent murine cell line L929 are shown in panels G and H [33].

By performing another experiment, we investigated the methionine dependence of the MM cell lines. In principle, methionine is essential, but cells can usually regenerate methionine from the precursor Hcy [28]. Many tumour cells, however, lose the ability to generate methionine from Hcy [26,29,34]. In the experiment, we used three metabolites from the methionine cycle, namely, Hcy, homoserine (HSe) and SAM. In MPC11, none of the three metabolites achieved compensation over a period of 72 h (Fig 2B). In the two human cell lines L363 (Fig 2D) and KMS12-BM (Fig 2F), only Hcy had a slight compensatory effect, but the methionine level did not reach the level of the control. In a previous work, we demonstrated the potential for methionine regeneration in the L929 murine cell line [35]. The results are shown in Fig 2G and H to demonstrate the functionality of the assay and the compensation for MetR by Hcy.

### Methionine-based AAR is antiproliferative, not apoptotic

Since methionine is an essential amino acid, the complete absence of the amino acid over a long period inevitably means potential damage to cell integrity, which can also lead to cell death. Since we completely removed methionine from the medium in our experiments, the potential for cell death in the cell lines used was investigated over a period of 72 h. No significant cell death was observed in any of the cell lines. No significant increase in cell death under MetR was observed in any of the cell lines (Fig 3). Staurosporine, a competitive inhibitor of the binding of ATP to various kinases that can induce cell death or apoptosis at relatively high concentrations, was used as a positive control (death control). Compared with the death control, in which all the cells in all the cell lines died after 72 h, all but a few cells were alive under MetR.

**Fig 3.**
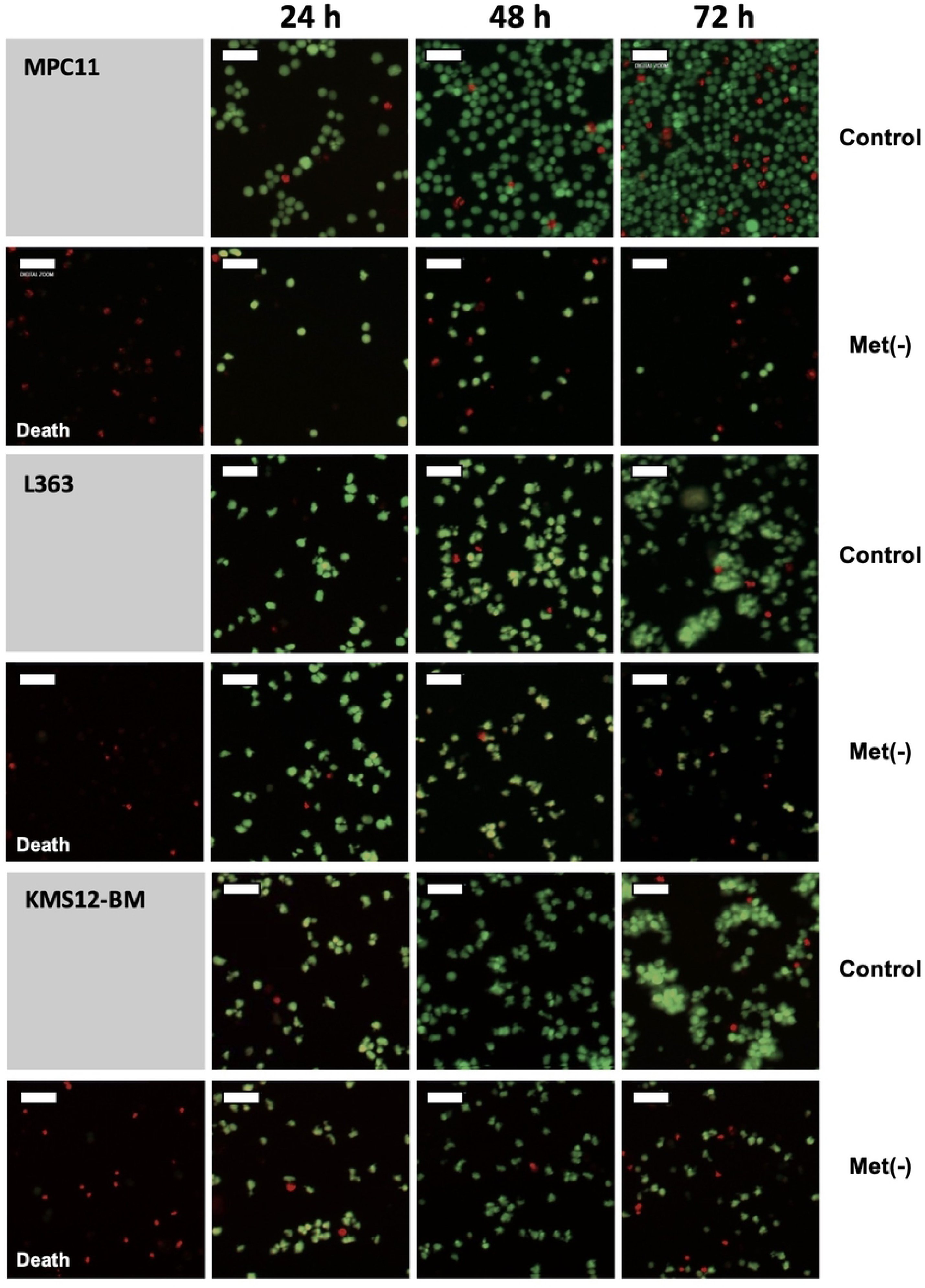
Analysis of Cell Viability. The cell lines MPC11, L363 and KMS12-BM were seeded at 10,000 cells/well in a 96-well plate. The cells were stimulated with control medium or Met(-)- medium. For the positive control (death), the cells were stimulated with 1 µM staurosporine. After 24 h, 48 h and 72 h, measurements were performed with the EarlyTox Live/Dead Assay Kit (Molecular Devices). The staining solution (5 µg/mL Hoechst 33342 (Thermo Fisher, Darmstadt, Germany), 6 µM EthD-III, and 6 µM calcein AM (CAM) in DMEM (Gibco, Life Technologies; Darmstadt, Germany)) was added, and after 30 min of incubation, measurements were made with the ImageXpress Pico Automated Cell Imaging System. The number of cells was determined via Hoechst 33342. Living cells were stained with the nonfluorescent calcein AM, which permeates the intact cell membrane and is converted into calcein, the fluorescent form, by intracellular esterases (cells are shown in green). If compromised cell membrane integrity is associated with cell death, EthD-III enters cells and binds to nucleic acids (cells are shown in red). A representative area for cell viability is shown for each cell line. The cell numbers are not representative of every case. The white bar, representing 52.44 µm, is used for size classification.

### LC‒MS analysis of MetR reveals metabolic reprogramming in MPC11 cells

We established the L929 murine cell line as a model system for analysing MetR via mass spectrometry [36]. Hence, we chose to use a murine cell line in this study for several reasons. First, much of the research on different types of nutrient restriction has been performed with rodent models. In this context, successful model systems for analysing energy metabolism have been established in mice [37]. Therefore, the use of a murine cell line allows better comparisons with results reported in the literature. Second, murine metabolism is considerably faster and more efficient than human metabolism. When the metabolic equivalents for energy/body weight/time (Kcal/g/h) are compared, the metabolic turnover of mice is as much as 100 times faster than that of humans [38–40]. For this reason, murine cells are much better suited than human cells for analyses of the metabolome. A third reason that we chose to analyse mouse cells was the many mechanisms that are highly conserved or triggered in very similar ways in mice and humans. With the beginning of the first cell, energy and mass are subjected to high selection pressure, which leads to the evolution of many basic mechanisms. Therefore, nutrient restriction studies have been performed in a wide range of organisms, from yeast to nematodes and from Drosophila and rodents to primates and even humans [19,21]. Many conserved mechanisms are evident, and although they have been adapted in individual species, these mechanisms are highly conserved. mTOR and sirtuins are only two examples of conserved mechanistic proteins [13,16]. The MPC11 murine cell line meets the conditions for use in the context of MM, as demonstrated by the experiments described herein. For these reasons, a murine cell line is also ideally suited to gain initial insights into metabolism under MetR via mass spectrometry.

Through LC‒MS, the influence of MetR on the metabolism of MPC11 cells was analysed every 24 h over a 5-day period. To ensure that as few cells as possible died or were under severe metabolic stress, a concentration of 0.5 mg/L methionine was used for these experiments under restrictive conditions (MetR). Only select results showing a clear trend over the entire study period are presented. No attention was given to the individual values at individual time points.

For most nonessential amino acids, lower concentrations and greater decreases in concentration compared with those of the control were observed up to the 120-h measurement (Fig 4). The opposite trend was observed for the essential amino acids, and the concentrations of these amino acids were maintained compared with the concentration of the control. The exceptions to this trend were the amino acids valine (Val), Gln, proline (Pro) and tyrosine (Tyr), which are shown in the lower part of the heatmap in Fig 4 and likely play a special role under MetR conditions. The amino acids Val, Gln and Tyr were stored in abundance under MetR conditions, and the amino acid Pro was similarly stored in abundance under control conditions. In addition, nutrient restriction clearly had a strong effect on the intracellular concentration of methionine (42% compared with the control) after only 24 h, and the methionine concentration continuously decreased, reaching a very low level (3%) over the course of the experiment.

**Fig 4.**
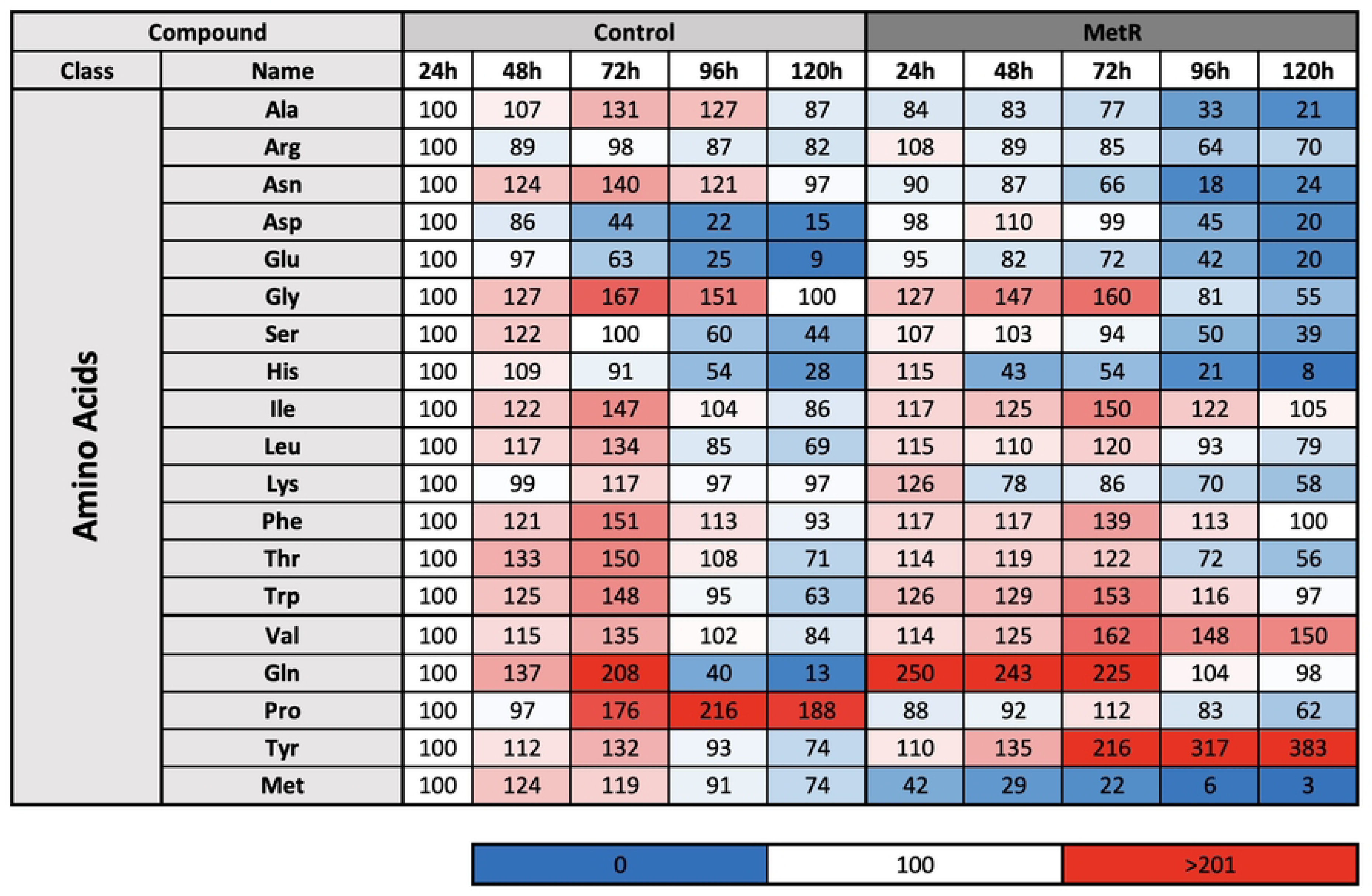
Relative Levels of Proteinogenic Amino Acids in MPC11 Cells Under Methionine Restriction (MetR) Conditions. The metabolism of this murine cell line was analysed in complete medium and MetR under proliferative conditions over a five-day period. For each day and condition of the experiment, triplicate samples were prepared. After 24 h, 48 h, 72 h, 96 h and 120 h, the cell lysates (intracellular) were analysed via liquid chromatography–mass spectrometry (LC‒MS). The results are reported as a summary of three independent experiments. For each analysis, the control value at 24 h was defined as 100%.

The concentrations of various polyamines, including spermidines and spermine, can reflect the energy level of a cell. Low intracellular concentrations can lead to the inhibition of protein synthesis and growth [41]. Similar effects can be achieved by the ingestion of extracellular spermidine by cells, leading to cell growth inhibition and autophagy induction [42,43]. In the case of MPC11 cells, spermidine and spermine do not exhibit a time course concentration that qualifies them as marker molecules for the cell energy state (Fig 5). The time course and concentration trends under MetR conditions were quite similar, except for a statistical outlier at 72 h. Creatine and creatinine play important roles in energy balance. Creatine phosphate is primarily an energy buffer in muscle and can phosphorylate ADP to form ATP when energy is rapidly needed. In the recovery phase, creatine is primarily rephosphorylated via a membrane-borne mitochondrial creatine kinase. Creatinine is the product of degraded creatine [44]. While the creatinine concentration was quite similar under both AAR conditions, the requirement under the MetR condition declined sharply after 96 h, reflecting a lower energy demand.

**Fig 5.**
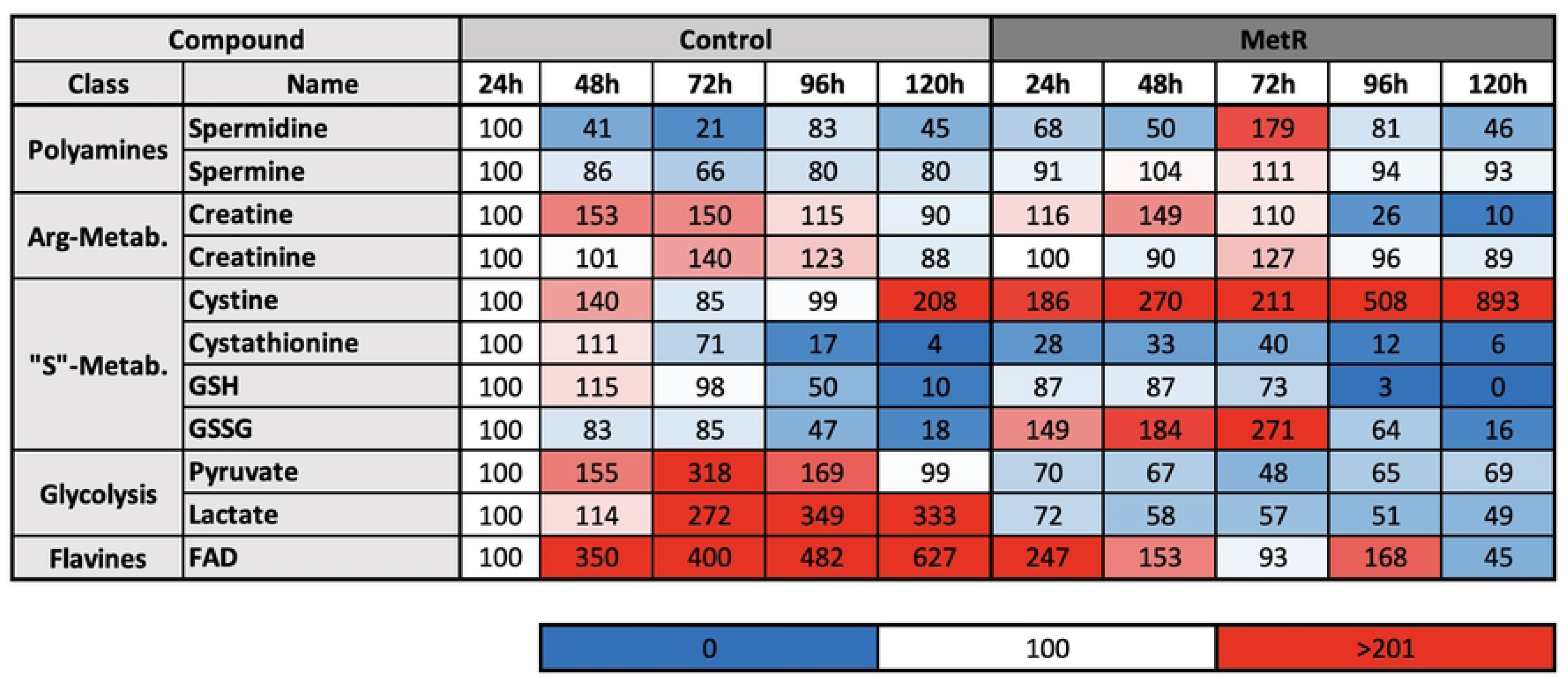
Polyamines, Arginine, Sulfur-Based and Metabolites of Glycolysis and Flavins in MPC11 Cells Under MetR. The metabolism of the murine cell line was analysed in complete medium under MetR and proliferative conditions over a five-day period. For each day and condition of the experiment, triplicate samples were prepared. After 24 h, 48 h, 72 h, 96 h and 120 h, the cell lysates (intracellular) were analysed via liquid chromatography–mass spectrometry (LC‒MS). The results are reported as a summary of three independent experiments. For analysis, the control value after 24 h was defined as 100%.

Sulfhydryl-based metabolism (“S”-Metab.) was among the most notable changes. One reaction to the absence of the sulphur source of methionine under the MetR condition was increased storage of the amino acid cysteine in the form of the disulfide cystine. Therefore, the concentration of the cysteine precursor cystathionine was lower than that of the control over the entire period under MetR conditions. The large storage requirement for cystine can possibly be explained by the high demand of haematopoietic cells for glutathione (GSH), which is formed from the three amino acids glutamate, cysteine and glycine. Gln, an amino acid that is also stored in increasing abundance in MPC11 cells under MetR conditions (Fig 4), is a central transport amino acid that is highly important in glutaminolysis, among other functions [45]. The antioxidant GSH is present at high concentrations in all cells but has additional significance in the stress reactions of leukocytes [46]. Under MetR conditions, the level of the oxidized form of GSH (GSSG) likely also profoundly increased.

Additionally, glycolysis significantly influenced the flow rate of metabolites. Pyruvate and lactate were found at profoundly reduced concentrations. The lower pyruvate concentration suggests a reduced glycolysis rate. Lactate, on the other hand, is a measure of the degree of cell proliferation, among other functions, specifically indicating inhibited cell proliferation. Increased lactate levels are not typical in neoplastic cells because of the Warburg effect, but they are generally higher in proliferating cells [47,48].

### MetR induced LEM and reduced nucleotide metabolism

Inhibiting cell proliferation is among the most prominent effects of LEM. In this context, nucleotides play a special role. Nucleotides form both the backbones of energy equivalents such as ATP and GTP and the backbones of RNA and DNA. LEM in MPC11 cells was characterized by a substantial reduction in the levels of many nucleotide metabolites, ranging from purines to pyrimidines (Fig 6). This reduction in nucleotide metabolites was explained by a lower demand for energy equivalents and by a reduced DNA synthesis rate due to inhibited cell proliferation. The effect on inhibited cell proliferation was particularly clear in the analysis of the metabolite carbamoyl aspartate, the level of which was much greater in the highly proliferative control group than in the inhibited proliferative group (a multiple-fold difference). Carbamoyl aspartate is a primer for pyrimidine biosynthesis.

**Fig 6.**
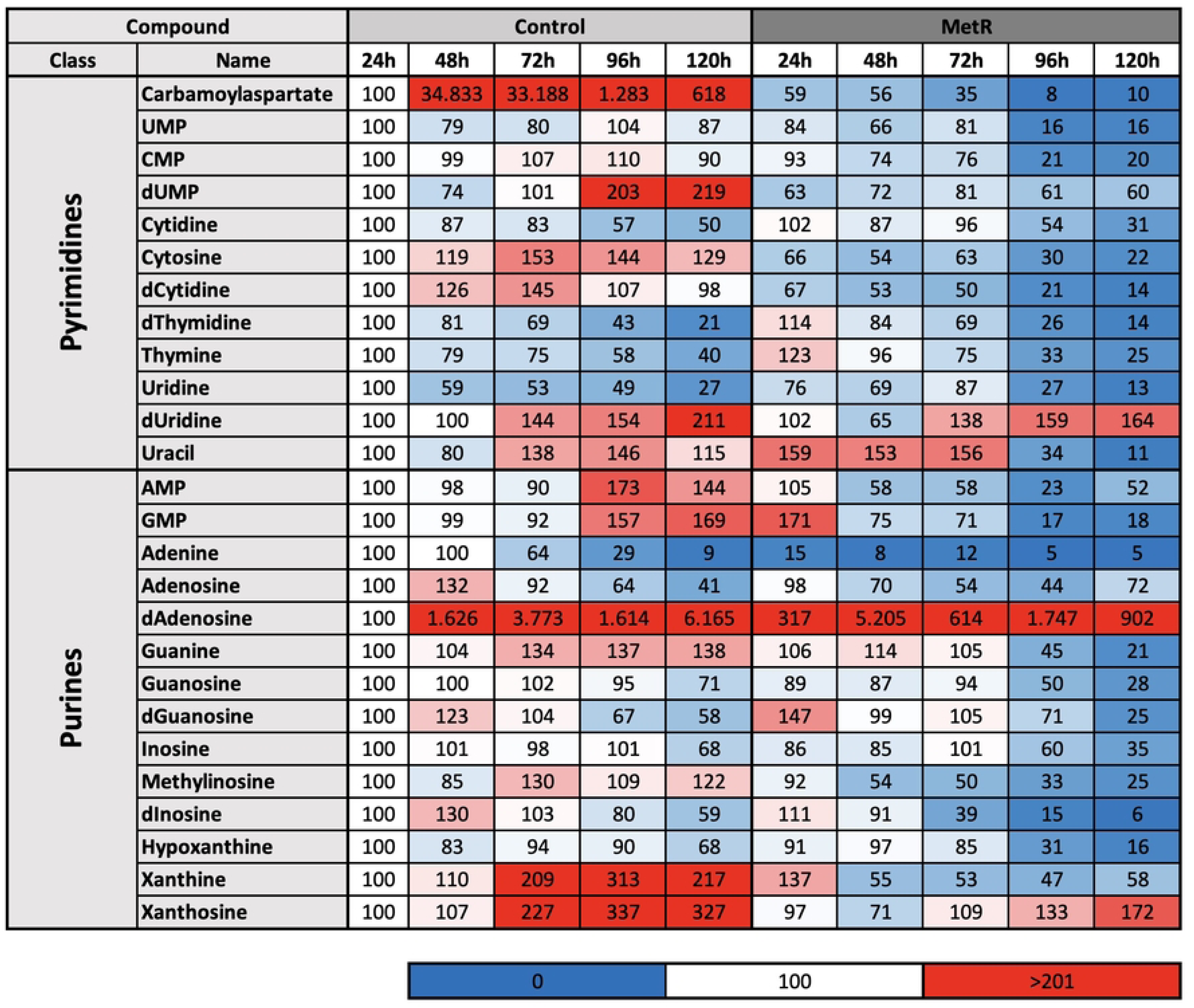
Relative Levels of Pyrimidines and Purines in MPC11 Cells Under Methionine Restriction (MetR) Conditions. The metabolism of this murine cell line was analysed in complete medium under MetR and proliferative conditions over a five-day period. For each day and condition of the experiment, triplicate samples were prepared. After 24 h, 48 h, 72 h, 96 h and 120 h, the cell lysates (intracellular) were analysed via liquid chromatography–mass spectrometry (LC‒MS). The results are reported as a summary of three independent experiments. For the analysis, the control value after 24 h was defined as 100%.

## Discussion

In summary, MetR was found to be a very successful nutrient restriction approach for inhibiting murine and human MM cell line proliferation. Cell methionine dependence is also highly important in potential tumour therapy, since Hcy cannot compensate for the lack of the amino acid methionine. All three cell lines were found to be methionine dependent. LC–MS analyses confirmed that MetR induced metabolic reprogramming in MPC11 cells. Notably, a reduced glycolysis rate and a dramatically reduced lactate level suggested a reduction in aerobic glycolysis (Warburg effect), as well as a continual reduction in the metabolites of energy equivalents and nucleotides obtained from groups of purines and pyrimidines.

Although we are the first to demonstrate that MetR efficiently suppresses murine and human MM cell line proliferation, the nutrient restriction approach is certainly not new. Notably, AAR constitutes an ideal approach in tumour therapy, which is the reason that the roles of amino acids have been highlighted again within the past decade [49,50]. The AAR approach is particularly suitable for haematological pathologies, as target cells can be effectively reached via the blood, especially as the blood, as both a carrier and a medium, contains numerous metabolites involved in key steps of amino acid metabolism.

In addition to AAR, LEM can be implemented via enzymatic depletion or, generally, through metabolic inhibition [51]. The effects of nutrient restriction methods, including CR, PR or AAR, on MM are largely unknown. Ohno and Cardullo [52] published “Effect of caloric restriction on neoplasm growth” in 1983, in which they reported the effectiveness of CR in treating leukaemia, among other diseases. In 1993, Mukkpadhyay et al. [53] published a comparative study showing the influence of CR on the growth of murine lymphoma. CR delays and inhibits tumour growth and prolongs host survival [53]. In another paper, a transgenic mouse model of B-cell lymphoma demonstrated that CR affected Mcl-1 expression and sensitized cells to BH3 mimetic-induced apoptosis independently of major BH3-only proteins and p53. This strategy may increase the efficiency of cancer therapy or prevent resistance to treatment [54].

Another possibility to reach the goal of depleting an amino acid involves conversion through (mostly) microbial enzymes that specifically degrade an amino acid and can often be simply administered through the oral route [55,56]. One clinically utilized strategy targeting amino acid dependency involves the use of asparaginase in the treatment of acute lymphoblastic leukaemia (ALL), which results in the depletion of exogenous asparagine and subsequent cancer cell death [51]. MetR, realized through the activity of methionase, has been investigated in phase I clinical trials involving patients with lymphoma and has led to a successful reduction in the serum methionine level [57].

Inhibition of membrane-localized amino acid transporters of the solute carrier (SLC) family or key intracellular enzymes involved in amino acid metabolism is possible. To achieve inhibitory effects in both cases, Gln is a typical target because of its central position in tumour metabolism [45]. Telaglenastat can inhibit glutaminase, a glutaminolysis enzyme that converts Gln to glutamate, enabling replenishment of the citrate cycle through the anaplerotic response [58]. The inhibition of key functions in MM cells by targeting Gln has been described [51]. V9302 is an inhibitor of SLC1A5, a specific membrane L-Gln transporter that can sensitize plasma myeloma cells to proteasome inhibitors [59]. With respect to the MM cells investigated in our study, the inhibition of membrane-localized cystine transporters is a very good approach that has been generally implemented in tumour therapy [60]. A nutrient restriction approach based on amino acid metabolism, as we describe herein, which is very useful in principle and has been previously described in the field of haematological pathologies, is far from being an established treatment. Although MetR is a useful approach because cells depend on methionine, the feasibility of using these approaches is not simple. Removing methionine from the diet or limiting its level is difficult. However, the aforementioned enzymes that degrade amino acids and inhibit amino acid transporters are therapies that can be effectively implemented.

Moreover, the spectrum of possible therapies can be expanded when MetR is included in the overall approach. As described in the Introduction, AAR represents only one possible form of nutrient restriction (with PR and CR being other forms) that can ultimately be implemented by leveraging a variety of molecular mechanisms that are common to all forms of nutrient restriction. MetR thus represents only one possible approach, but in this work, we demonstrated the efficiency with which MetR can inhibit cell proliferation. In addition, the therapy spectrum can be expanded by CRMs, which constitute a class of substances with mechanisms of action that are not precisely defined and are extremely heterogeneous. In summary, any approach that can produce the same effect as CR, PR or AAR can be defined as a CRM. We define the overall effect of nutrient restriction and CRM as “metabolic silencing”, i.e., the switching of metabolism to LEM through different approaches or agents separately or in combination.

The methionine dependence of MM cells is high, which can be demonstrated by the use of methionine radiotracers in positron emission tomography (PET)/computed tomography (CT) in MM, which enables a much more accurate diagnosis [61], as glucose-based methods often do not guarantee sufficient imaging owing to hexokinase-2 deficiency in MM cells [62]. In principle, MetR can also be induced indirectly by drugs already used in the treatment of MM. Immunomodulatory drugs such as lenalidomide may be able to indirectly induce a form of MetR, as the application of the drug leads to a reduction in the aforementioned amino acid transporter SLC7A5 (LAT1), which transports methionine, among other things [63]. The effect of the drug can therefore be attributed, among other things, to a reduction in amino acids.

In recent years, metformin, which is a prime example of a CRM, has played a special role in treating disease. Metformin has a variety of effects. For example, the energy sensor AMPK is activated via the inhibition of mitochondrial electron transport chain complex I [64,65]. Alternatively, metformin can directly inhibit mTOR independently of the action of AMPK [66]. Metformin can increase deacetylation of the proteome, e.g., by increasing the activity of the NAD+-dependent histone deacetylase sirtuin 1 [67]. Metformin is an attractive agent in MM therapy. For example, metformin has been shown to inhibit IL-6 signalling by decreasing IL-6R expression in MM cells [68]. The efficacy of metformin against haematological diseases has also been demonstrated in numerous studies, but no clinical studies have been performed to date to demonstrate its efficacy [69,70].

Another advantage of “metabolic silencing” is its ability to induce an extreme condition called “differential stress resistance”. Cancer cells that express oncogenes and exhibit egocentric proliferative behaviour respond to certain cancer-promoting and growth-promoting factors. Moreover, cancer cells do not respond to protective signals generated by short-term CR (SCR) or long-term nutrient restriction. Hence, cells can be exposed to two different extreme situations: somatic cells may be protected, while cancer cells become increasingly vulnerable to attack [71]. This condition can be achieved, for example, by SCR. The first SCR studies were performed with patients diagnosed with diffuse large B-cell lymphoma. SCR was associated with improvements in postchemotherapy haematological parameters (i.e., erythrocyte and lymphocyte counts) [72].

The increased vulnerability of tumour cells to “differential stress resistance” was also shown through LC–MS analyses. In our work, we focused on amino acids under MetR conditions, and notably, Tyr (Fig 4) was found to accumulate in cells under MetR conditions. Two interpretations can explain this finding: either Tyr consumption was greater under MetR conditions, and the cells had higher Tyr demand, or Tyr was stored for later use. Since cells under MetR conditions stop proliferating, the latter interpretation probably offers a better explanation. Cells that proliferate rapidly need much more Tyr than cells with LEM. Therefore, an inhibitor of Tyr would profoundly affect proliferating cells but only slightly affect cells with LEM.

In summary, the results of this work indicate that MetR is a very efficient approach for inhibiting MM cell proliferation. In addition, the efficiency of these approaches is supported by the dependence of the cell lines studied on the extracellular supply of methionine. Considering the transferability of the results to human MM cell lines, metabolic silencing via methionine-based AAR is a potential strategy for MM therapy.

## Supporting Information

**S1 Data Tables Fig 1. Analysis of the Cell Proliferation Rate.** (PRISM file)

**S2 Data Tables Fig 2. Analysis of Methionine Resistance (MetR) and Homocysteine (Hcy) S3 Data Table Mass Spectrometry Methionine Restriction L929.** (Excel file)

**Supporting information captions**

## Acknowledgments

This publication was supported by the Open Access Publication Fund of the University of Wuerzburg.

## Author Contributions

Conceptualization: Axel Seher

Investigation: Andrea Frabschka, Maximilian F. Völter, Hannah Jöhren, Werner Schmitz, Charlotte Wiegand, Mohamed El-Mesery, Lena Wiesner

Methodology: Corinna Koderer Supervision: Axel Seher

Writing – original draft: Axel Seher

Writing – review & editing: Alexander Christian Kübler

## References

1. Rajkumar SV. Multiple myeloma: 2011 update on diagnosis, risk-stratification, and management. Am J Hematol. 2011;86: 57–65.

2. Longo V, Brunetti O, D’Oronzo S, Dammacco F, Silvestris F. Therapeutic approaches to myeloma bone disease: an evolving story. Cancer Treat Rep. 2012;38: 787–797.

3. Palumbo A, Anderson K. Multiple myeloma. N Engl J Med. 2011;364: 1046–1060.

4. Harousseau JL, Moreau P. Autologous hematopoietic stem-cell transplantation for multiple myeloma. N Engl J Med. 2009;360: 2645–2654.

5. Rajkumar SV. Multiple myeloma: every year a new standard? Hematol Oncol. 2019;37: 62–65.

6. Gahvari Z, Brunner M, Schmidt T, Callander NS. Update on the current and future use of CAR-T to treat multiple myeloma. Eur J Haematol. 2024;112: 493–503.

7. Kumar SK, Dispenzieri A, Lacy MQ, Gertz MA, Buadi FK, Pandey S, et al. Continued improvement in survival in multiple myeloma: changes in early mortality and outcomes in older patients. Leukemia. 2014;28: 1122–1128.

8. NIH. Cancer stat facts: myeloma. 2022 [cited 14 Febrauary 2022]. Available from: https://seer.cancer.gov/statfacts/html/mulmy.html.

9. Landgren O, Iskander K. Modern multiple myeloma therapy: deep, sustained treatment response and good clinical outcomes. J Intern Med. 2017;281: 365–382.

10. Hanahan D, Weinberg RA. Hallmarks of cancer: the next generation. Cell. 2011;144: 646–674.

11. Hosios AM, Hecht VC, Danai LV, Johnson MO, Rathmell JC, Steinhauser ML, et al. Amino acids rather than glucose account for the majority of cell mass in proliferating mammalian cells. Dev Cell. 2016;36: 540–549.

12. Gongol B, Sari I, Bryant T, Rosete G, Marin T. AMPK: an epigenetic landscape modulator. Int J Mol Sci. 2018;19: 3238.

13. Longo VD, Kennedy BK. Sirtuins in aging and age-related disease. Cell. 2006;126: 257–268.

14. Goberdhan DC, Wilson C, Harris AL. Amino acid sensing by mTORC1: intracellular transporters mark the spot. Cell Metab. 2016;23: 580–589.

15. Lauinger L, Kaiser P. Sensing and signaling of methionine metabolism. Metabolites. 2021;11: 83.

16. Kim J, Guan KL. mTOR as a central hub of nutrient signalling and cell growth. Nat Cell Biol. 2019;21: 63–71.

17. Escobar KA, Cole NH, Mermier CM, VanDusseldorp TA. Autophagy and aging: maintaining the proteome through exercise and caloric restriction. Aging Cell. 2019;18: e12876.

18. Alves-Fernandes DK, Jasiulionis MG. The role of SIRT1 on DNA damage response and epigenetic alterations in cancer. Int J Mol Sci. 2019;20: 3153.

19. Ball ZB, Barnes RH, Visscher MB. The effects of dietary caloric restriction on maturity and senescence, with particular reference to fertility and longevity. Am J Physiol. 1947;150: 511–519.

20. Gillespie ZE, Pickering J, Eskiw CH. Better living through chemistry: caloric restriction (CR) and CR mimetics alter genome function to promote increased health and lifespan. Front Genet. 2016;7: 142.

21. Mirzaei H, Suarez JA, Longo VD. Protein and amino acid restriction, aging and disease: from yeast to humans. Trends Endocrinol Metab. 2014;25: 558–566.

22. Madeo F, Carmona-Gutierrez D, Hofer SJ, Kroemer G. Caloric restriction mimetics against age-associated disease: targets, mechanisms, and therapeutic potential. Cell Metab. 2019;29: 592–610.

23. Harris RE, Beebe-Donk J, Doss H, Doss DB. Aspirin, ibuprofen, and other non-steroidal anti-inflammatory drugs in cancer prevention: a critical review of non-selective COX-2 blockade (review). Oncol Rep. 2005;13: 559–583.

24. Foretz M, Guigas B, Bertrand L, Pollak M, Viollet B. Metformin: from mechanisms of action to therapies. Cell Metab. 2014;20: 953–966.

25. Ables GP, Hens JR, Nichenametla SN. Methionine restriction beyond life-span extension. Ann N Y Acad Sci. 2016;1363: 68–79.

26. Cavuoto P, Fenech MF. A review of methionine dependency and the role of methionine restriction in cancer growth control and life-span extension. Cancer Treat Rep. 2012;38: 726–736.

27. Wanders D, Hobson K, Ji X. Methionine restriction and cancer biology. Nutrients. 2020;12: 684.

28. Finkelstein JD. Methionine metabolism in mammals. J Nutr Biochem. 1990;1: 228–237.

29. Borrego SL, Lin DW, Kaiser P. Isolation and characterization of methionine-independent clones from methionine-dependent cancer cells. Methods Mol Biol. 2019;1866: 37–48.

30. Kreis W. Tumor therapy by deprivation of L-methionine: rationale and results. Cancer Treat Rep. 1979;63: 1069–1072.

31. Gubser PM, Kallies A. Methio “mine”! cancer cells steal methionine and impair CD8 T-cell function. Immunol Cell Biol. 2020;98: 623–625.

32. Bian Y, Li W, Kremer DM, Sajjakulnukit P, Li S, Crespo J, et al. Cancer SLC43A2 alters T cell methionine metabolism and histone methylation. Nature. 2020;585: 277–282.

33. Wünsch AC, Ries E, Heinzelmann S, Frabschka A, Wagner PC, Rauch T, et al. Metabolic silencing via methionine-based amino acid restriction in head and neck cancer. Curr Issues Mol Biol. 2023;45: 4557–4573.

34. Chaturvedi S, Hoffman RM, Bertino JR. Exploiting methionine restriction for cancer treatment. Biochem Pharmacol. 2018;154: 170–173.

35. Schmitz W, Ries E, Koderer C, Völter MF, Wünsch AC, El-Mesery M, et al. Cysteine restriction in murine L929 fibroblasts as an alternative strategy to methionine restriction in cancer therapy. Int J Mol Sci. 2021;22: 11630.

36. Schmitz W, Koderer C, El-Mesery M, Gubik S, Sampers R, Straub A, et al. Metabolic fingerprinting of murine L929 fibroblasts as a cell-based tumour suppressor model system for methionine restriction. Int J Mol Sci. 2021;22: 3039.

37. Azzu V, Valencak TG. Energy metabolism and ageing in the mouse: a mini-review. Gerontology. 2017;63: 327–336.

38. Melzer K, Heydenreich J, Schutz Y, Renaud A, Kayser B, Mäder U. Metabolic equivalent in adolescents, active adults and pregnant women. Nutrients. 2016;8: 438.

39. National Research Council Subcommittee on Laboratory Animal Nutrition. Nutrient requirements of laboratory animals. Washington, DC: National Academies Press; 1995.

40. Speakman JR. Body size, energy metabolism and lifespan. J Exp Biol. 2005;208: 1717–1730.

41. Mandal S, Mandal A, Johansson HE, Orjalo AV, Park MH. Depletion of cellular polyamines, spermidine and spermine, causes a total arrest in translation and growth in mammalian cells. Proc Natl Acad Sci U S A. 2013;110: 2169–2174.

42. Chen Y, Zhuang H, Chen X, Shi Z, Wang X. Spermidine–induced growth inhibition and apoptosis via autophagic activation in cervical cancer. Oncol Rep. 2018;39: 2845–2854.

43. Madeo F, Eisenberg T, Pietrocola F, Kroemer G. Spermidine in health and disease. Science. 2018;359: eaan2788.

44. Wallimann T, Tokarska-Schlattner M, Schlattner U. The creatine kinase system and pleiotropic effects of creatine. Amino Acids. 2011;40: 1271–1296.

45. Hensley CT, Wasti AT, DeBerardinis RJ. Glutamine and cancer: cell biology, physiology, and clinical opportunities. J Clin Invest. 2013;123: 3678–3684.

46. Muri J, Kopf M. Redox regulation of immunometabolism. Nat Rev Immunol. 2021;21: 363–381.

47. Liberti MV, Locasale JW. The warburg effect: how does it benefit cancer cells? Trends Biochem Sci. 2016;41: 211–218.

48. Vander Heiden MG, Cantley LC, Thompson CB. Understanding the Warburg effect: the metabolic requirements of cell proliferation. Science. 2009;324: 1029–1033.

49. Bastings J, Van Eijk HM, Damink SWO, Rensen SS. d-amino acids in health and disease: a focus on cancer. Nutrients. 2019;11: 2205.

50. Butler M, Van Der Meer LT, Van Leeuwen FN. Amino acid depletion therapies: starving cancer cells to death. Trends Endocrinol Metab. 2021;32: 367–381.

51. Endicott M, Jones M, Hull J. Amino acid metabolism as a therapeutic target in cancer: a review. Amino Acids. 2021;53: 1169–1179.

52. Ohno T, Cardullo AC. Effect of caloric restriction on neoplasm growth. Mt Sinai J Med. 1983;50: 338–342.

53. Mukhopadhyay P, Gupta JD, Sanyal U, Das S, Senyal U. Influence of dietary restriction and soyabean supplementation on the growth of a murine lymphoma and host immune function. Cancer Lett. 1994;78: 151–157.

54. Meynet O, Zunino B, Happo L, Pradelli LA, Chiche J, Jacquin MA, et al. Caloric restriction modulates Mcl-1 expression and sensitizes lymphomas to BH3 mimetic in mice. Blood. 2013;122: 2402–2411.

55. Dhankhar R, Gupta V, Kumar S, Kapoor RK, Gulati P. Microbial enzymes for deprivation of amino acid metabolism in malignant cells: biological strategy for cancer treatment. Appl Microbiol Biotechnol. 2020;104: 2857–2869.

56. Wang Z, Xie Q, Zhou H, Zhang M, Shen J, Ju D. Amino acid degrading enzymes and autophagy in cancer therapy. Front Pharmacol. 2020;11: 582587.

57. Tan Y, Zavala J, Han Q, Xu M, Sun X, Tan X, et al. Recombinant methioninase infusion reduces the biochemical endpoint of serum methionine with minimal toxicity in high-stage cancer patients. Anticancer Res. 1997;17: 3857–3860.

58. Cederkvist H, Kolan SS, Wik JA, Sener Z, Skålhegg BS. Identification and characterization of a novel glutaminase inhibitor. FEBS Open Bio. 2022;12: 163–174.

59. Prelowska MK, Mehlich D, Ugurlu MT, Kedzierska H, Cwiek A, Kosnik A, et al. Inhibition of the ʟ-glutamine transporter ASCT2 sensitizes plasma cell myeloma cells to proteasome inhibitors. Cancer Lett. 2021;507: 13–25.

60. Koppula P, Zhuang L, Gan B. Cystine transporter SLC7A11/xCT in cancer: ferroptosis, nutrient dependency, and cancer therapy. Protein Cell. 2020;12: 599–620.

61. Lapa C, Kircher M, Da Via M, Schreder M, Rasche L, Kortüm KM, et al. Comparison of 11C-choline and 11C-methionine PET/CT in multiple myeloma. Clin Nucl Med. 2019;44: 620–624.

62. Rasche L, Angtuaco E, McDonald JE, Buros A, Stein C, Pawlyn C, et al. Low expression of hexokinase-2 is associated with false-negative FDG-positron emission tomography in multiple myeloma. Blood. 2017;130: 30–34.

63. Heider M, Eichner R, Stroh J, Morath V, Kuisl A, Zecha J, et al. The IMiD target CRBN determines HSP90 activity toward transmembrane proteins essential in multiple myeloma. Mol Cell. 2021;81: 1170–1186.e10.

64. Duca FA, Côté CD, Rasmussen BA, Zadeh-Tahmasebi M, Rutter GA, Filippi BM, et al. Metformin activates a duodenal Ampk-dependent pathway to lower hepatic glucose production in rats. Nat Med. 2015;21: 506–511.

65. Owen MR, Doran E, Halestrap AP. Evidence that metformin exerts its anti-diabetic effects through inhibition of complex 1 of the mitochondrial respiratory chain. Biochem J. 2000;348: 607–614.

66. Nair V, Sreevalsan S, Basha R, Abdelrahim M, Abudayyeh A, Hoffman AR, et al. Mechanism of metformin-dependent inhibition of mammalian target of rapamycin (mTOR) and Ras activity in pancreatic cancer: role of specificity protein (Sp) transcription factors. J Biol Chem. 2014;289: 27692–27701.

67. Caton PW, Nayuni NK, Kieswich J, Khan NQ, Yaqoob MM, Corder R. Metformin suppresses hepatic gluconeogenesis through induction of SIRT1 and GCN5. J Endocrinol. 2010;205: 97–106.

68. Mishra AK, Dingli D. Metformin inhibits IL-6 signaling by decreasing IL-6R expression on multiple myeloma cells. Leukemia. 2019;33: 2695–2709.

69. Podhorecka M. Metformin -its anti-cancer effects in hematologic malignancies. Oncol Rev. 2021;15: 514.

70. Cunha AD, Pericole FV, Carvalheira JBC. Metformin and blood cancers. Clinics (Sao Paulo). 2018;73: e412s.

71. Lee C, Longo VD. Fasting vs dietary restriction in cellular protection and cancer treatment: from model organisms to patients. Oncogene. 2011;30: 3305–3316.

72. Tang CC, Huang TC, Tien FM, Lin JM, Yeh YC, Lee CY. Safety, feasibility, and effects of short-term calorie reduction during induction chemotherapy in patients with diffuse large B-cell lymphoma: a pilot study. Nutrients. 2021;13: 3268.

